# Characterization of Chromium-Resistant and Reducing Bacteria from Poultry Litter Ecosystems

**DOI:** 10.64898/2026.06.30.734947

**Authors:** Tamanna Zerin, Maria Islam Bethe, Sumaiya Sultana, Sabrina Aktar, Marufa Akter, Md. Abdullah Ibna Masud, Sardar Muhammad Osail

## Abstract

Compact poultry raising has turned poultry litter into an environmental problem, as it may all be packed with heavy metals and drug-resistant germs. Of all the metals, chromium contamination not only disturbs the general environment but is also a source of concern for public health. Poultry litters were taken from 14 farms in different places, and the bacteria characters from different places were tested for their capacity to tolerate Cr(VI). A total of 31 bacterial isolates were initially screened, and three of them (AH-2, AZ-1, and AMF-3) appeared to be very resistant to chromium. The isolates were able to survive at the highest concentration, 800 mg/L of the Cr(VI); however, AH-2 was the most resistant one (MIC: 900 mg/L; MBC: 1000 mg/L). Chromium reduction tests showed that AMF-3 at high concentration showed the maximum chromium reduction, while AH-2 achieved higher chromium reduction at medium concentration. Phenotypic and biochemical analysis showed that the isolates were *Staphylococcus* spp., which was confirmed by 16S rRNA gene sequencing as *S. cohnii, S. saprophyticus*, and *S. gallinarum*. Moreover, chromium was detected at higher levels in poultry litter compared to the feed, with the highest accumulation in AZ farm litter (4464.0 μg/kg). The highlighting feature of our article is the presence of chromium-tolerant and reducing bacteria in poultry environments. Besides that, the level of chromium in poultry litter is really high, and it points to the need for better waste management.

## Introduction

Over the past centuries, industrialization has been expanding indiscriminately without considering the disposal process of effluents. The expulsion of heavy metal-containing solid or liquid wastes from various industries, including leather tanning, textile, and electroplating, among others, causes environmental pollution with metals. Among them, hexavalent chromium, Cr(VI), has augmented the great concern about human health. Among different oxidation states, Cr(III) and Cr(VI) are the most abundant chromium compounds in the natural environment, with different physicochemical characteristics and distinct biological interactions. Of these two states, trivalent chromium is less stable and relatively insoluble and immobile than its hexavalent form (Khan et al. 2024; Shahriar et al. 2025). Research showed that the hexavalent form of Cr is a carcinogenic agent for humans and 500 times more toxic than the trivalent one (Shahriar et al. 2025; Nur-E-Alam et al. 2020). The presence of toxic heavy metals in the human food chain may cause damage to human organs like bone marrow, kidney, and liver, as well as cause dermatitis, gastrointestinal ulcers, and cancers (Bokhtiar et al. 2023). Because of the high demand for poultry meat, the poultry industry is rapidly growing in developing countries to meet the demand. Side by side, poultry embodies a threat to human health, specifically as a vector of infectious diseases and due to the presence of chromium-resistant isolates (Mottet and Tempio 2017). Nowadays, tannery waste has become a good source for animal feed, and there are around 70 big and medium-sized and 300 small feed factories in Bangladesh that prefer tannery waste for animal feed (Wilbur et al. 2012). A study on the discharge of tannery solid waste showed that various tannery mills and many locals were involved in converting the solid waste into protein-concentrate for mixing into poultry feed. Around 200-250 tons of protein concentrate are produced per day, which is then turned into feed and sold to fish and poultry feed factories (Mazumder et al. 2013). Another study in Bangladesh, conducted by Dhaka University and Bangladesh Council for Scientific and Industrial Research (BCSIR), revealed higher amounts (3.2037 %) of chromium in eggs and poultry meat than the tolerable limit (Mazumder et al 2013), followed by 2.4901% as an element of protein concentrate sampled from a feed mill in Hazaribagh. Due to improper disposal and recycling practices, tannery waste is notorious for its high Cr content and often finds its way into animal feed, including chicken feed, further exacerbating Cr accumulation in chickens (Akter et al. 2024). Several studies showed the correlation between chromium tolerance and MDR property due to its co-selection pressure of antibiotics and presence of chromium reductase enzyme (Durjoy et al. 2025; Jain et al. 2012). However, researchers are interested to use enzymatic property from chromium tolerant bacteria for their potential in bioremediation. This study is focused on isolating and characterizing chromium-tolerant bacteria from poultry litter as well as determining resistance potential and the ability of these bacteria to reduce chromium. Another objective of the study is to measure chromium contamination in poultry feed and litter.

## Materials and Method

### Study Area and Sample Collection

The research was conducted between February and April 2025 with 14 poultry farms across three different districts of Bangladesh, namely Dhaka, Sirajganj, and Narayanganj (**Fig 1**). To identify and maintain records, every poultry farm was assigned an abbreviation.

**Fig 1.**
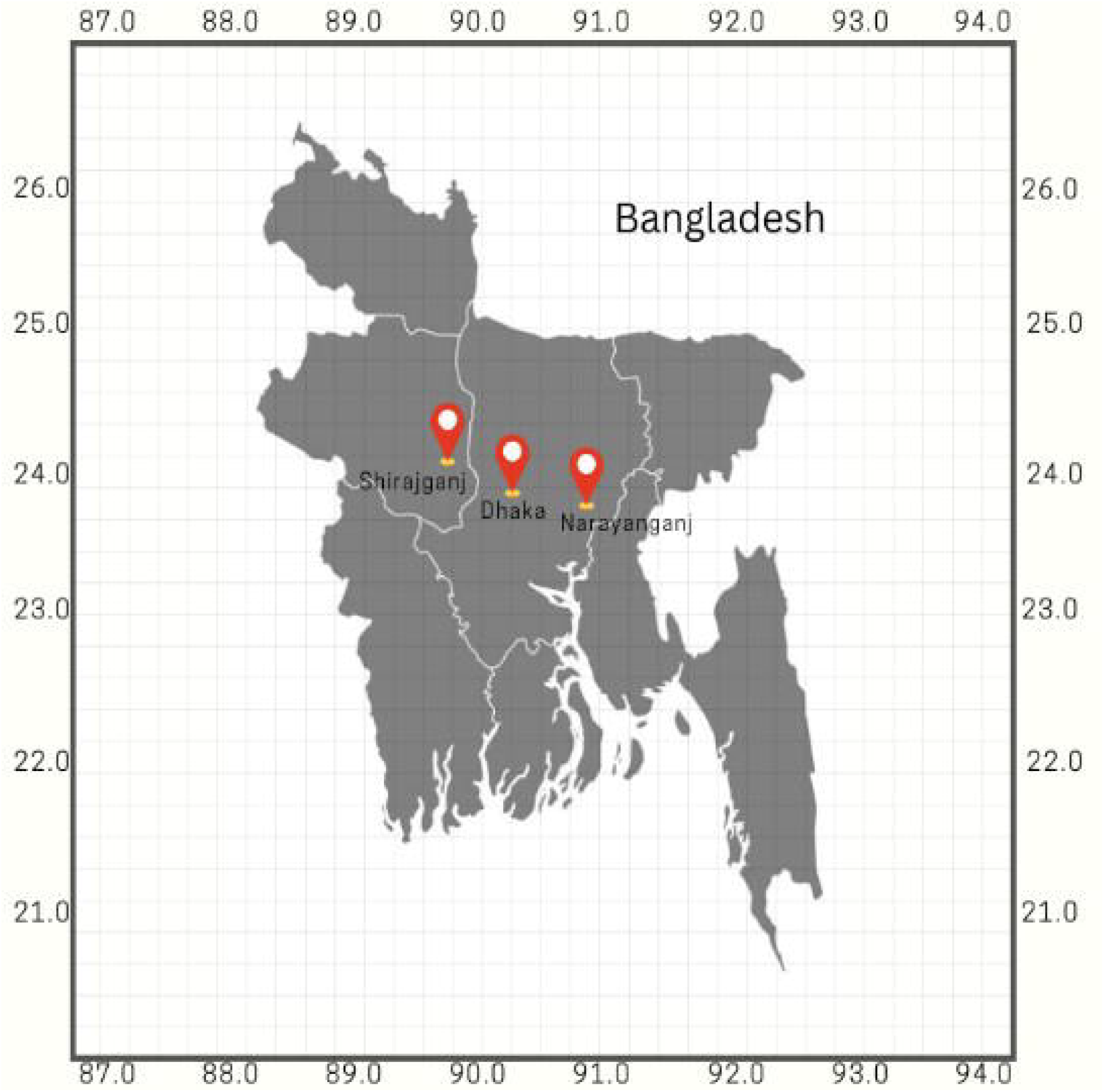
Sample collection area

Samples of feed and litter were taken with great attention to hygiene in the exposure to infection under outdoor conditions at various times of the day. Litter samples were obtained from the various areas of the poultry house (e.g., corners, center, and near feeders and drinkers) using sterile spatulas to achieve representativeness, and then, all the samples were mixed to create a composite sample. Around 50-100 g of litter was put in sterile, marked polyethylene bags.

Feed samples were directly taken from feeding trays or storage containers in a sterile way. About 50 g of feed from each farm was aseptically transferred into sterile containers. Every sample was properly marked, then taken in insulated boxes to the laboratory, and finally, they were immediately dealt with for the subsequent microbiological testing.

### Sample Processing, Isolation, and Screening of Bacteria

The collected feed and litter samples were collected in a sterile bag and stored at 4°C. Within 24 hours, samples were measured and serially diluted in sterile normal saline from 10□¹ to 10□□and then distributed in aliquots from dilutions 10□³ to 10□□ onto nutritional agar (HIMEDIA, India) plates supplemented with 100 mg/L potassium dichromate (K_2_Cr_2_O_7_) using the spread plate technique. Inoculated plates were incubated at 37°C for 24 hours. Selected single colonies were streaked onto NA plates to obtain a pure culture of isolates. Isolated colonies were used for further identification of colony characteristics, microscopic traits, and biochemical tests.

### Determination of Cr(VI) Tolerance

Chromium tolerance was assessed by culturing isolates in NA media containing 200–1100 mg/L Cr(VI), prepared from a 125,000 mg/L K□Cr□O□stock solution. Cultures were incubated at 37°C for 24 hours, and growth was evaluated visually. Isolates growing at ≥800 mg/L were classified as highly resistant. Tolerance was scored semi-quantitatively as heavy, moderate, slight, and no growth.

### Determination of Minimum Inhibitory Concentration (MIC) and Minimum Bactericidal Concentration (MBC)

The minimum inhibitory concentration (MIC) and minimum bactericidal concentration (MBC) of hexavalent chromium, Cr(VI), were assessed by the broth microdilution method. Bacterial strains were grown in Luria Broth (LB) at 37°C for 24 h and adjusted to a standard inoculum equivalent to 0. 5 McFarland standard (around 1×10□CFU/mL). Two-fold dilutions of Cr(VI) were made in a sterile 96-well microtiter plate to get final concentrations from 100 to 3200 g/mL. Each well was inoculated with this bacterial suspension. Wells with only medium and Cr(VI) and wells with only bacteria were used as negative and positive controls, respectively. The plates were placed at 37°C for 24 h, and the MIC was recognized as the lowest concentration of Cr(VI) showing no visible bacterial growth compared to controls.

In determining the MBC, the wells with no visible growth in the MIC test were picked out, and colonies were identified on NA plates after incubation at 37°C for 24 h. The MBC represented the lowest level of Cr(VI), where no bacterial colonies appeared on the plates, indicating the total killing of bacteria.

### Chromium (VI) Reduction Assay

Based on tolerance screening, three highly resistant isolates (AH02, AZ01, AMF03) were selected for quantitative Cr(VI) reduction using the 1,5-diphenylcarbazide (DPC) method. Isolates were grown in LB broth with 1.56–200 mg/L Cr(VI) at 37°C for 24 h. Then, cultures were centrifuged, and the pH of supernatants was adjusted to 2.0 with 2 M H□SO□. DPC reagent (0.5 mL; 250 mg DPC in 50 mL acetone) was added, and the Cr(VI)-DPC complex was measured at 540 nm. Decreased color intensity relative to an uninoculated control indicated Cr(VI) reduction, and the percentage reduction was calculated as:

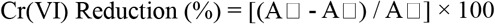

Where A□= Absorbance before incubation, A□= Absorbance after incubation.

### Presumptive Identification

To determine the bacterial isolates capable of tolerating extremely elevated concentrations of Cr(VI), first their morphology (e.g., whether they exhibited rod- or sphere-like structure) and nature of aggregation were examined, as well as their Gram staining (i.e., whether positive or negative). Following the selection of isolates, various biochemical tests were performed to confirm that these bacteria were capable of growth via these mechanisms (citrate utilization, triple sugar iron, oxidase and catalase tests), with each test containing positive and negative controls. To further aid in the identification of these isolates, selective media like mannitol salt agar (MSA) were also used.

### Molecular Characterization

For molecular identification, genomic DNA was extracted using the QIAamp DNA Mini Kit (QIAGEN, Germany). The 16S rRNA gene was amplified by PCR with universal primers, 27F (5’-AGAGTTTGATCMTGGCTCAG-3’) and 1492R (5’-GGTTACCTTGTTACGACTT-3’), and a clear amplicon of approximately 1500 bp was obtained. The products were purified with ExoSAP-IT (Thermo Fisher Scientific). Cycle sequencing was performed with the BigDye Terminator v3.1 Kit (Applied Biosystems), purified using a DyeEx 2.0 Spin Kit (QIAGEN), and run on an ABI 3500 XL Genetic Analyzer. The resulting 16S rRNA sequences were compared to GenBank using BLASTn (NCBI) to determine closest phylogenetic relatives and assign species identities.

### Quantification of Chromium in Poultry Feed and Litter

Total chromium in poultry feed and litter was determined by atomic absorption spectroscopy (AAS) after wet acid digestion. Approximately 0.4 g of each dried, homogenized sample was weighed into a digestion tube, mixed with 5 mL concentrated HNO□, and left to stand for 1 hour. Then, 3 mL HClO□was added, and the mixture was allowed to stand for 30 minutes. Samples were digested on a hot plate at 200°C until red nitrous fumes disappeared and the solution became clear. After cooling, the digest was diluted to 25 mL with deionized water and filtered through

Whatman No. 44 filter paper. Chromium concentration in the filtrate was measured using an atomic absorption spectrophotometer.

## Results

### Characteristics of Sampled Farms

During the period February-April 2025, a total of 14 poultry farms in three districts, namely Dhaka (Khilgaon, Shekhertake), Shirajganj (Janpur), and Narayanganj (Kachpur), were selected for sampling. In order to identify each farm, they were given unique abbreviations. Sampling was carried out at various times of the day in field conditions. Across all farms, the main bedding materials were wood powder and/or wood chaff, and the houses were bedded for 1 to 60 days before sampling. The differences in the age of bedding were used to see the possible effect on microbial and physicochemical characteristics. The farms obtained feed from various commercial sources: Teer Company, Nahar Agro, Pronita Company, CP feed, Kazi feed, Pustiraj feed, and RRP feed, besides one farm which used combined layer feed and cow feed. These different farm management practices, like the kind of bedding, duration, and feed source, were factored for the purpose of representative sampling that will allow the assessment of environmental and microbial issues in poultry production systems.

**Table 01:** Location and temporal details of chicken litter sample collection.

| Farm Name | Abbreviation | Location | Date | Time | Bedding Type | Bedding Duration | Feed |
| --- | --- | --- | --- | --- | --- | --- | --- |
| Mamun Enterprise | ME | Khilgaon, Dhaka | 11.02.2025 | 11.02 pm | Wood powder, chaff | 10 days older | Teer Company |
| Nusaiba Poultry Farm | NF | Shekhertake, Dhaka | 17.02.2025 | 12.10 pm | Wood powder, chaff | 15 days older | Nahar Agro |
| Personal farm | PF | Shekhertake, Dhaka | 17.02.2025 | 12.25 pm | Wood powder, chaff | 27 days older | Pronita Company |
| Alamin Farm 01, | AF 01 | Janpur, Shirajganj | 22.02.2025 | 4.50 pm | Wood powder | 1 day older | CP feed |
| Alamin Farm 02, | AF 02 | Janpur, Shirajganj | 22.02.2025 | 5.00 pm | Wood Powder | 30 days older | CP feed |
| Hasan Agro | HA | Janpur, Shirajganj | 05.04.2025 | 5.50 pm | Wood chaff | 12 days older | Kazi feed |
| Abdul Monem Farm | AMF | Janpur, Shirajganj | 05.04.2025 | 6.00 pm | Wood chaff | 60 days older | Layer-1+ cow feed |
| Tania Farm | TF | Janpur, Shirajganj | 05.04.2025 | 5.10 pm | Wood chaff | 4 days older | Pustiraj feed |
| Mamun Farm | MF | Janpur, Shirajganj | 05.04.2025 | 5.05 pm | Wood chaff | 45 days older | Pustiraj feed |
| Jahidul Farm | JF | Kachpur, Narayanganj | 09.04.2025 | 3.28 pm | Wood powder, chaff | 16 days older | RRP feed |
| Hamim Farm 01 | HM 01 | Kachpur, Narayanganj | 09.04.2025 | 3.40 pm | Wood powder, chaff | 1 day older | RRP feed |
| Hamim Farm 02 | HM 02 | Kachpur, Narayanganj | 09.04.2025 | 3.50 pm | Wood powder, chaff | 30 days older | RRP feed |
| Azizul Farm | AZ | Kachpur, Narayanganj | 09.04.2025 | 4.10 pm | Wood powder, chaff | 25 days older | RRP feed |
| Abed Hasan Farm | AH | Kachpur, Narayanganj | 09.04.2025 | 4.30 pm | Wood powder, chaff | 12 days older | RRP feed |

### Isolation of the Highest Chromium-Tolerant Bacteria

A total of 31 bacterial isolates were recovered from 15 poultry litter samples collected from farms in Dhaka and surrounding areas, and screened for 100 mg/L Cr(VI) tolerance. Among these, three isolates (AH-2, AZ-1, and AMF-3) exhibited the highest tolerance to hexavalent chromium. These isolates originated from litter samples of Abed Hasan Farm (Kachpur), Azizul Farm (Kachpur), and Abdul Monem Farm (Shirajganj).

Chromium tolerance assays demonstrated substantial variation in bacterial growth across Cr(VI) concentrations ranging from 200 to 1100 mg/L. Isolates AH-2 and AZ-1 maintained heavy growth up to 800 mg/L, while AMF-3 exhibited heavy growth up to 600 mg/L and moderate growth at higher concentrations. Growth decreased progressively as chromium concentration increased, and no growth was observed at 1100 mg/L for all isolates.

### Minimum Inhibitory and Bactericidal Concentrations (MIC and MBC)

Comparison of the minimum inhibitory concentration (MIC) and minimum bactericidal concentration (MBC) results showed that all isolates followed the same trend. The lowest MIC (700 mg/L) and MBC (900 mg/L) were recorded for the isolate AMF-3, which would be more sensitive to chromium exposure than the other strains. On the other hand, AZ-1 had the next level of resistance with MIC and MBC being 800 mg/L and 1000 mg/L, respectively. The highest level of tolerance was exhibited by AH-2 with a MIC of 900 mg/L and an MBC of 1000 mg/L. Interestingly, the MBC levels for all strains were always higher than their corresponding MICs, confirming that the bactericidal doses prevail over the inhibitory concentrations. Besides, this behavior is parallel with a dose-dependent effect and reveals differences in the level of chromium resistance among the strains, namely, AH-2, the most resistant, and AMF-3, the most sensitive (**Fig 2**).

**Fig 2.**
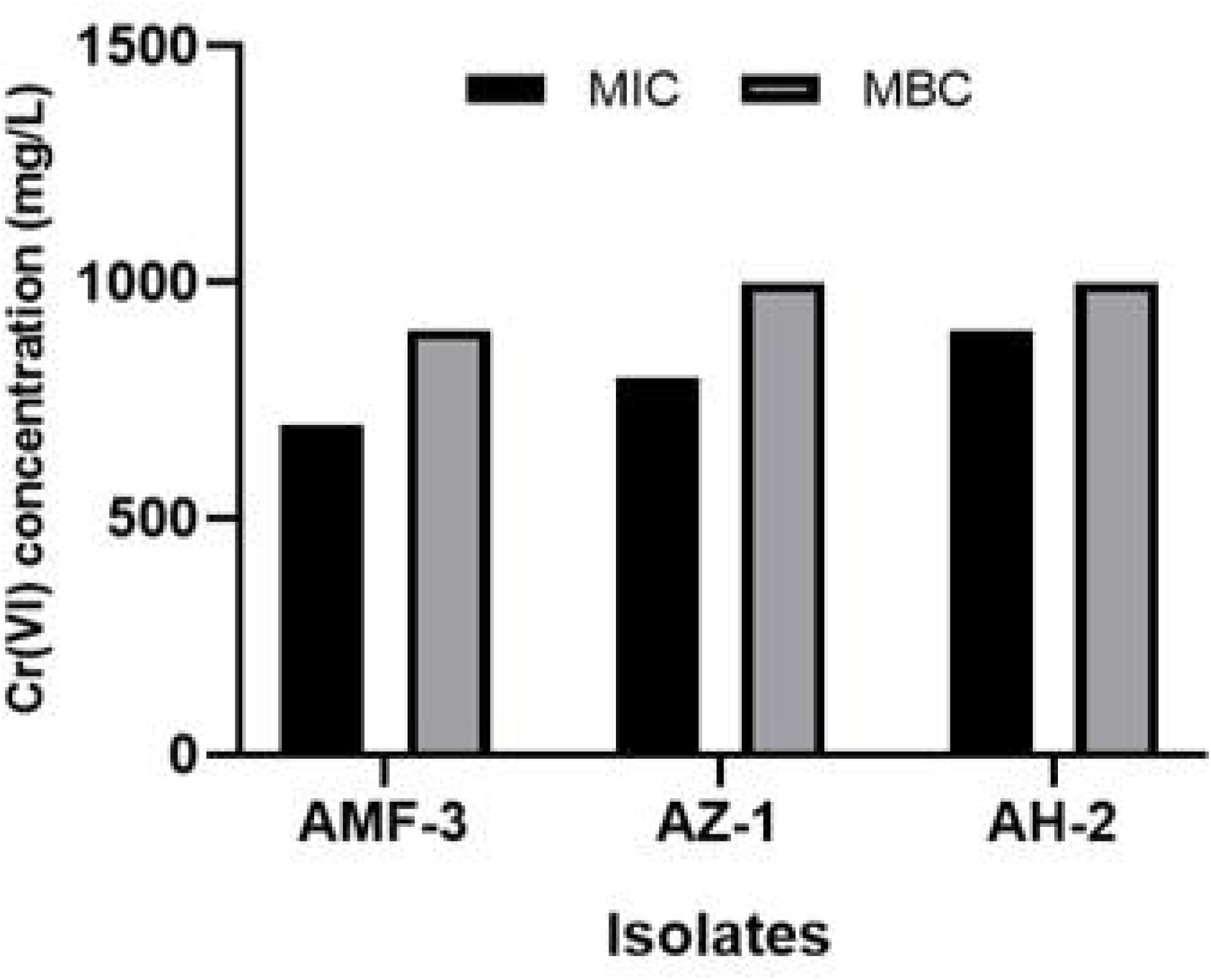
MIC and MBC of the highest Cr(VI) tolerant isolates

**Table 2:** Cr(VI) tolerance profile of the selected isolates at 200 to 1100 mg/L concentrations.

| ID | Cr(IV) in mg/L |  |  |  |  |  |  |  |  |  |
| --- | --- | --- | --- | --- | --- | --- | --- | --- | --- | --- |
|  | 200 | 300 | 400 | 500 | 600 | 700 | 800 | 900 | 1000 | 1100 |
| ME-01 | +++ | +++ | +++ | ++ | ++ | + | - | - | - | - |
| ME-02 | +++ | +++ | +++ | + | + | + | - | - | - | - |
| ME-03 | +++ | +++ | +++ | +++ | + | + | - | - | - | - |
| ME-04 | +++ | +++ | +++ | ++ | + | + | - | - | - | - |
| ME-05 | +++ | +++ | +++ | +++ | +++ | +++ | + | - | - | - |
| NF-01 | +++ | +++ | +++ | + | + | - | - | - | - | - |
| NF-02 | +++ | +++ | +++ | - | - | - | - | - | - | - |
| NF-03 | +++ | +++ | +++ | + | - | - | - | - | - | - |
| PF-01 | +++ | +++ | +++ | ++ | + | + | - | - | - | - |
| PF-02 | +++ | +++ | +++ | + | + | + | - | - | - | - |
| PF-03 | +++ | +++ | +++ | ++ | + | + | - | - | - | - |
| AF01-01 | +++ | +++ | +++ | + | + | + | - | - | - | - |
| AF01-02 | +++ | +++ | +++ | + | + | + | - | - | - | - |
| AF02-01 | +++ | +++ | +++ | ++ | - | - | - | - | - | - |
| AF02-02 | +++ | +++ | +++ | + | + | - | - | - | - | - |
| AMF-01 | +++ | +++ | +++ | +++ | +++ | +++ | + | - | - | - |
| AMF-02 | +++ | +++ | +++ | ++ | + | - | - | - | - | - |
| **AMF-03** | +++ | +++ | +++ | ++ | +++ | + | + | + | - | - |
| MF-01 | +++ | +++ | +++ | - | - | - | - | - | - | - |
| MF-02 | +++ | +++ | +++ | + | - | - | - | - | - | - |
| TF-01 | +++ | +++ | +++ | +++ | ++ | - | - | - | - | - |
| TF-02 | +++ | +++ | +++ | +++ | + | + | + | - | - | - |
| HA-01 | +++ | +++ | +++ | ++ | + | + | + | - | - | - |
| JF-01 | +++ | +++ | +++ | +++ | ++ | - | - | - | - | - |
| **AZ-01** | +++ | +++ | +++ | +++ | +++ | +++ | +++ | + | + | - |
| AZ-02 | +++ | +++ | +++ | +++ | ++ | ++ | + | - | - | - |
| AH-01 | +++ | +++ | +++ | +++ | ++ | + | - | - | - | - |
| **AH-02** | +++ | +++ | +++ | +++ | +++ | +++ | +++ | ++ | + | - |
| HM01-01 | +++ | +++ | +++ | +++ | +++ | ++ | - | - | - | - |
| HM01-02 | +++ | +++ | +++ | + | - | - | - | - | - | - |
| HM02-01 | +++ | +++ | +++ | +++ | ++ | - | - | - | - | - |

note: heavy growth (+++), moderate growth (++), slight growth (+), and no growth (-).

### Chromium Reduction Potential

The chromium reduction potentials of the three isolates (AH-2, AZ-1, and AMF-3) differed markedly across Cr concentrations, highlighting differences in both the efficiency of individual isolates and their responses to Cr concentration. AMF-3 topped the list of reduction efficiency at 200 mg/L (84.52%), followed by AZ-1 (35.50%) and AH-2 (25.09%), which also means AMF-3 had the best tolerance and reduction ability at high chromium levels. AMF-3 still had the highest reduction potential of about 89.46% at 100 mg/L, but AH-2 performed better (81.87%) than AZ-1 (54.71%). At moderate Cr levels (50-25 mg/L), AH-2 showed the highest reduction ability (nearly 83.49% and 86.93%, respectively), beating AMF-3 and AZ-1 and indicating that it is most active at intermediate Cr concentrations. When the Cr levels were very low (6.25-12.5 mg/L), AMF-3 still had the highest reduction ability (40.05% and 46.61%, respectively), while AZ-1 and AH-2 were only moderately effective. The activity of the isolates was almost nil at the lowest concentrations of 3.125-1.5625 mg/L, with AZ-1 showing a nearly negligible activity even at 1. 5625 mg/L. In fact, AMF-3 was the only one that showed the most consistent and highest reduction throughout the wide concentration range, especially at high Cr stress. AH-2 was the most efficient at moderate concentrations, and AZ-1 maintained a comparatively low reduction potential throughout the whole experiment (**Fig 3**).

**Fig 3.**
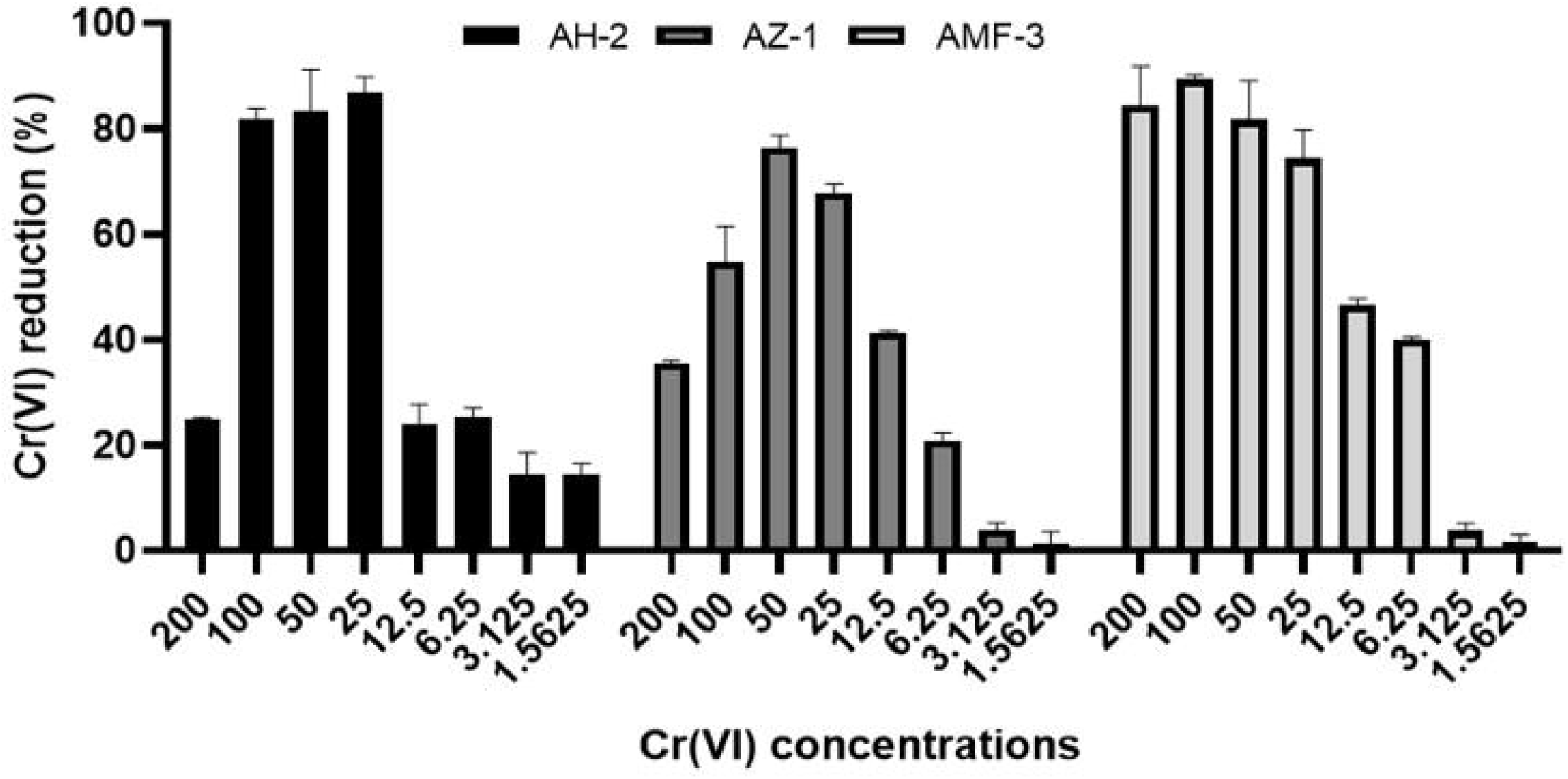
Cr(VI) reduction potential of the selected isolates at different Cr concentrations

### Phenotypic Characterization

The presumptive characterization of the isolates based on their cultural, microscopic, and biochemical features suggested that they were very similar to *Staphylococcus* spp. Growth on Mannitol Salt Agar (MSA) exhibited typical morphology in agreement with *Staphylococcus aureus*. Microscopic observation showed Gram-positive cocci arranged in clusters, which is the typical characteristic of the *Staphylococcus* genus. Biochemical analyses strongly confirmed this identification as these isolates were catalase positive, oxidase negative, and unable to use citrate. In the Triple Sugar Iron (TSI) test, all isolates turned the slant and the butt alkaline (K/K) without the formation of gas or hydrogen sulfide. Together, these phenotypic and biochemical features indicate that the isolates can be tentatively considered *Staphylococcus* spp.

**Table 3:** Phenotypic characterization of the selected isolates.

| Isolates ID | Gram staining | MSA media | Catalase | Oxidase | Citrate | TSI |  |  |  | Presumptive Identification |
| --- | --- | --- | --- | --- | --- | --- | --- | --- | --- | --- |
|  |  |  |  |  |  | Slant | Butt | H2O | Gas |  |
| AZ-01 | Cocci, single, pair, cluster, Gram + | Small, red, without yellow halo | + | - | - | K | K | - | - | *Staphylococcus* spp. |
| AH-02 | Cocci, single, pair, cluster, Gram + | Small, red, without yellow halo | + | - | - | K | K | - | - | *Staphylococcus* spp. |
| AMF-03 | Cocci, single, pair, cluster, Gram + | Small, white, with yellow halo | + | - | - | K | K | - | - | *Staphylococcus* spp. |

note: mannitol salt agar (msa), triple sugar iron (tsi), alkaline (k).

### Molecular Identification

Phylogenetic analysis of the 16S rRNA gene sequences clearly showed that these three isolates are taxonomically placed within the genus *Staphylococcus*. The phylogenetic tree that we made had very high bootstrap support (99%) for all major nodes, which means the clustering was very strong. Isolate AZ-1 was very close to *Staphylococcus cohnii*, and together with the reference strains including *S. cohnii* subsp. urealyticus, they made a well-supported clade. On the other hand, isolate AMF-3 belongs to the *Staphylococcus saprophyticus* group and is very similar to several reference strains such as J34, 55QC2CO, and MF B8, which is strongly supported by a 99% bootstrap value. Isolate AH-2 however is a member of the *Staphylococcus gallinarum* group and is very similar to strains like C25, SSB39, and 77QC2O2, again with strong bootstrap support (99%). The low evolutionary distance (scale bar: 0.002 substitutions per nucleotide position) suggests that the sequences of the isolates and their respective reference strains are very similar to one another, which is further evidence that their species-level identification is correct (**Fig 4**).

**Fig 4.**
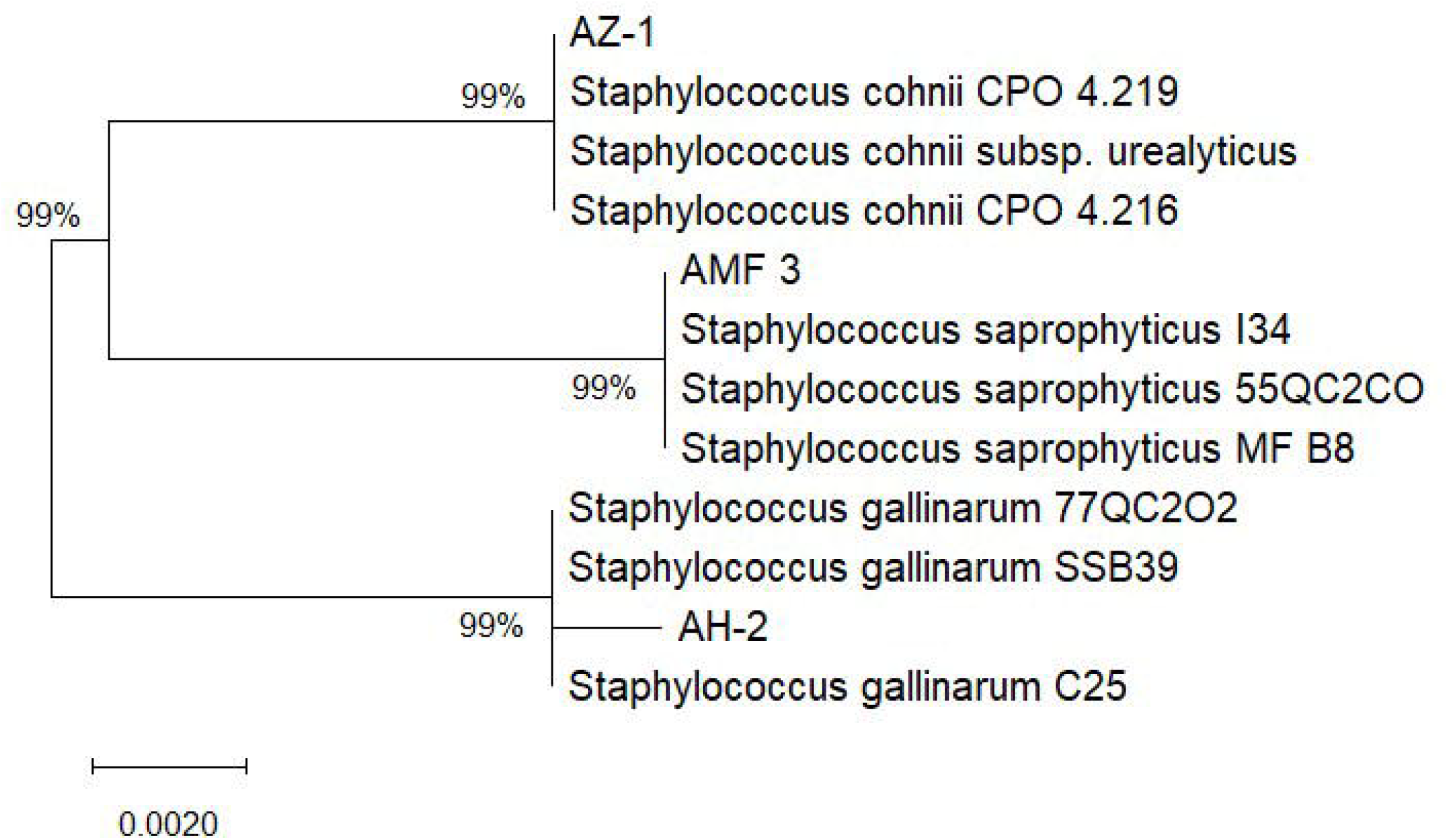
The phylogenetic tree was drawn from 16S rRNA gene sequences for the isolates AZ-1, AMF-3, and AH-2. The isolates correspond to *Staphylococcus cohnii, S. saprophyticus*, and *S. gallinarum* in their clusters, respectively. Bootstrap values (%) are displayed at nodes, and 0.002 substitutions per site is the length of the scale bar

### Chromium Concentration in Poultry Feed and Litter

Chromium contents in chicken feed and litter samples from the three farms (AH, AZ, and AMF) differed. The highest chromium concentration in the feed sample was that of farm AMF (2146.0 μg/kg), followed by AH (856.0 μg/kg). The lowest concentration of chromium was found in AZ (687.0 μg/kg). However, chromium levels in chicken litter were much higher than those in feed for all farms, with AZ having the highest concentration (4464.0 μg/kg), followed by AMF (1923.0 μg/kg) and AH (1385.0 μg/kg). Between the farms, AMF had the highest chromium load in feed, while AZ had the highest accumulation in litter, which shows different levels of chromium input and waste handling practices among the farms.

**Table 4:** Detection of Cr concentrations in chicken feed and litter at three different farms where the highest Cr(VI) tolerant bacterial isolates were identified.

| Types of samples | Chromium concentrations (µg/Kg) in farms |  |  |
| --- | --- | --- | --- |
|  | AH | AZ | AMF |
| Chicken feed | 856.0 | 687.0 | 2146.0 |
| Chicken litter | 1385.0 | 4464.0 | 1923.0 |

## Discussion

This paper offers an extensive examination of chromium pollution, microbial changes, and the ability to clean the environment using bacteria associated with poultry droppings from some farms in Bangladesh. Differences were found in the farm’s features, such as floor material, litter usage duration, and feed type, which conform to earlier studies showing that how poultry are managed affects microbial variation and the accumulation of heavy metals in litter (Oyewale et al. 2019; Bolan et al. 2004). The usage of wood-related floor materials and older litter age (up to 60 days) might have led to increased microbial count and growth in metal-absorbing ability, since it has been demonstrated that older litter accumulates higher levels of heavy metals due to repeated defecation and limited removal (Naveenkumar et al. 2026).

A total of 31 bacterial isolates were recovered, of which AH-2, AZ-1, and AMF-3 stood out as highly resistant to Cr(VI). The fact that these isolates could keep on growing at very high levels of chromium (up to 800 mg/L) is in line with previous findings that environmental bacteria, especially those coming from contaminated or waste rich environments, can become resistant through various adaptive mechanisms such as efflux systems, enzymatic reduction and metal sequestration (Cheung and Gu, 2007; Yu et al. 2023). Decrease in growth with increasing Cr(VI) concentration in this work is a typical dose-dependent inhibitory effect of chromium toxicity, as has been reported in resistance studies with some other microbes (Plestenjak et al. 2022).

The MIC and MBC results revealed different levels of resistance among the isolates. AH-2 was the most resistant (MIC: 900 mg/L; MBC: 1000 mg/L), while AMF-3 was the most sensitive. Typically, MBC values are higher than

MIC values, as obviously the killing doses are higher than the inhibitory ones (Andrews, 2001). Similar results were obtained from chromium-resistant *Staphylococcus* spp. and some other Gram-positive bacteria isolated from industrial and agricultural environments (Shakoori et al. 1999). The differences in resistance can stem from the genetic makeup, plasmid-borne resistance, or metabolic profiles of each isolate.

The chromium reduction assay highlighted clear functional variances among the isolates. AMF-3 showed the most effective reduction at high chromium levels, which implies a powerful enzymatic chromium reduction ability, likely through chromate reductases. This level of chromium reduction efficiency has also been found in *Staphylococcus saprophyticus* and other bacteria, which have the ability to convert Cr(VI) to Cr(III), the less toxic form (Pushkar et al. 2021). AH-2, on the other hand, exhibited better reduction at moderate levels, which means that the reduction efficiency might rely not only on the resistance level but also on the best metabolic conditions. The relatively poorer performance of AZ-1 points towards some strain-specific limitations in reduction mechanisms. These results confirm the observations of several other authors pointing out that chromium reduction depends on the strain as well as the concentration (Mishra et al. 2012).

The phenotypic and biochemical tests revealed that the isolates were from the genus *Staphylococcus*, and 16S rRNA gene sequencing confirmed it. The identification of isolates as *Staphylococcus cohnii, S. saprophyticus*, and *S. gallinarum* is very interesting, as these members of this genus are mainly associated with the animal environment and some members have been reported to have metal tolerance (Stepanovi et al. 2003). Traditionally, *Staphylococcus* spp. is not considered the first choice for bioremediation agents. However, there are reports showing their heavy metal resistance and the ability to detoxify metal ions, particularly in waste-rich ecological niches (Maki and Al-Taee 2023). The high bootstrap values in phylogenetic analysis indicate the accuracy of the taxonomic placement and the close genetic relationship of the isolates to the reference strains.

The study of chromium levels in the poultry feed and litter demonstrated a definite pattern where the litter consistently contained higher amounts of chromium than the feed. This finding aligns well with earlier studies that have shown poultry tend to excrete a large fraction of the heavy metals they eat, which then get accumulated in the litter (Bolan et al. 2004; Islam et al. 2008). The very high chromium level in AZ litter (4464.0 μg/kg) suggests environmental hazards if the waste is not properly managed. The variations seen between different farms - for example, AMF having higher chromium in feed and AZ showing higher chromium accumulation in litter - probably result from differences in feed formulations, supplementation methods, and litter management systems. In general, chromium buildup was higher in litter than in feed, which may be attributed to the bioaccumulation and excretion of chromium by poultry. There have been similar reports in poultry production where the use of feed additives and mineral supplements results in heavy metal fluctuations (Naveenkumar et al. 2026).

The discovery of chromium-resistant and chromium-reducing *Staphylococcus* spp. opens up their possible use in clean-up methods through bioremediation. Nevertheless, more research into their genome, enzyme activities, and the performance of these bacteria at a large scale is required in order to uncover their advantages on a practical level. Besides that, the tighter control of feed components along with the upgrading of the poultry farm waste handling should be prioritized to prevent environmental contamination and the risk to the health of the general public.

## Conclusion

Our research points out a massive build-up of chromium in poultry litter and, at the same time, reveals some chromium-resistant *Staphylococcus* spp. that have great reduction capacity. Such results underscore their potential for bioremediation of the contaminated agro-environment. Going forward, molecular mechanisms and large-scale implementation should be studied to develop sustainable approaches to combat heavy metals.

## Statements and Declarations

### Competing interests

The authors have no competing interests to declare that are relevant to the content of this article

### Funding

Our study was partially funded by the Ministry of Science and Technology, Government of Bangladesh, under the Project ID: R&D-2410112.

## Notes

### Competing Interest Statement

The authors have declared no competing interest.

